# User-friendly transcriptomic data analysis with ArrayAnalysis

**DOI:** 10.64898/2026.07.13.738193

**Authors:** Jarno Koetsier, Ozan Cinar, Egon L. Willighagen, Ammar Ammar, Vishnu Karthik, Danyel Jennen, Chris T. Evelo, Leopold M.G. Curfs, Chris P. Reutelingsperger, Nasim Bahram Sangani, Lars M.T. Eijssen

## Abstract

Transcriptomic profiling has become a cornerstone of modern biomedical research. To make transcriptomic analyses accessible to a broader scientific community, specifically including researchers with limited bioinformatics expertise, we introduced ArrayAnalysis in 2013 as a user-friendly web-based application for microarray data analysis. We now present a major update (https://arrayanalysis.org), introducing a strongly interactive platform that facilitates the dedicated exploration and analysis of both microarray and RNA-seq data, and allows for the generation of publication-ready outputs. Users can perform key analysis steps, including data pre-processing and quality control, differential expression analysis, and gene set analysis, via a sequential, interactive workflow. At each step, the application provides interactive visualizations accompanied by information pages to support interpretation. Users can dynamically adjust figure layouts and colour palettes and export figures as vector graphics and high-resolution raster images. For non-expert users, ArrayAnalysis offers step-by-step guidance to support correct usage and facilitate learning, while for experienced bioinformaticians, it provides a streamlined and flexible workflow ideal for large-scale analyses requiring efficient and consistent processing. ArrayAnalysis is available both as a web application and for local deployment as a desktop application, Docker image, or R package, making it suitable for diverse computational environments, user groups, and analytical purposes. Together, ArrayAnalysis empowers a broad community of biomedical researchers to unlock the full potential of transcriptomic data.

**GRAPHICAL ABSTRACT:** 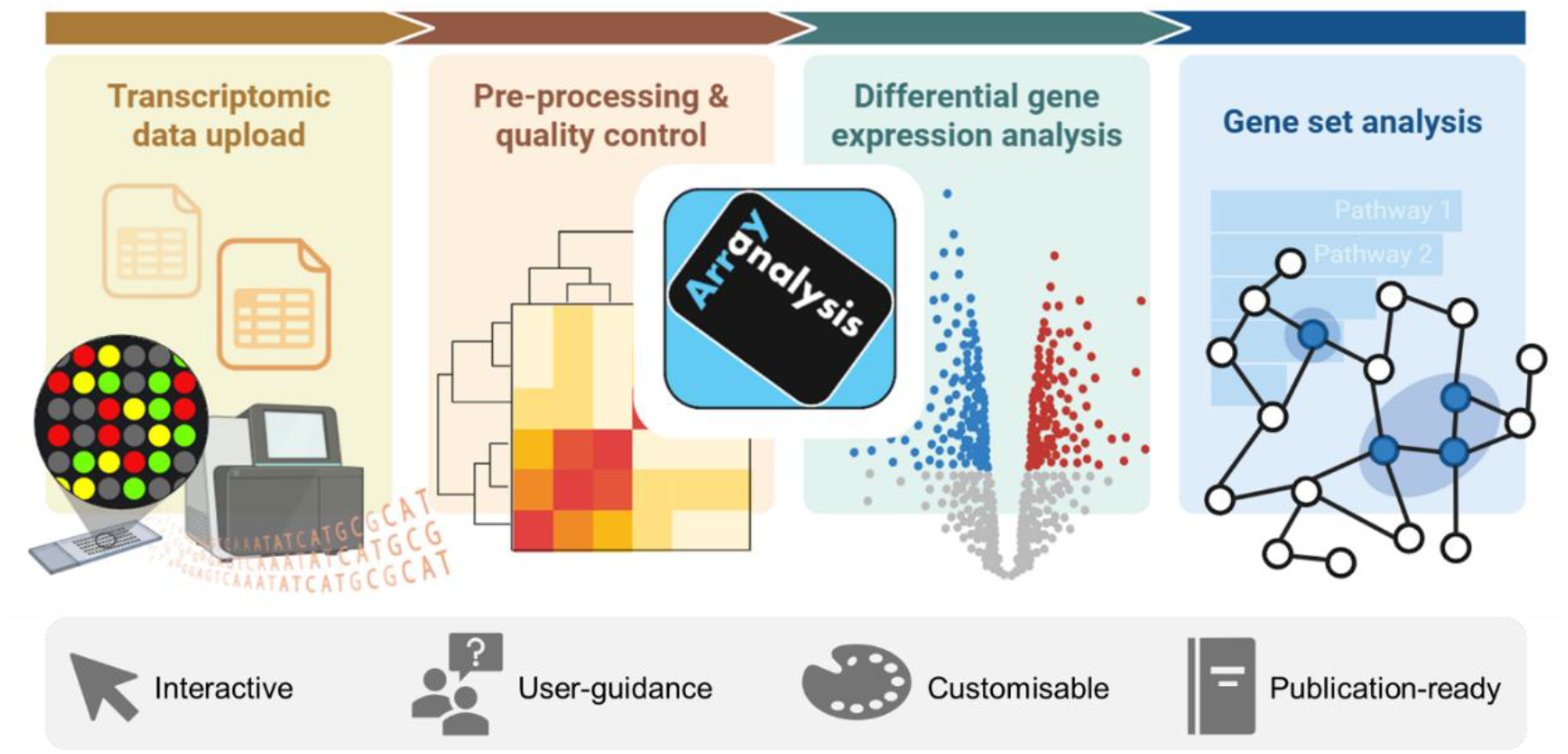

## BACKGROUND

Gene expression profiling, through microarray or RNA sequencing (RNA-seq) technologies, has become an indispensable tool in biomedical research for uncovering molecular mechanisms and identifying biomarkers. However, the analysis of transcriptomic data remains challenging for many researchers, especially for those without formal training in bioinformatics. To bridge this gap, several applications have been developed that offer a user-friendly interface to the analysis of transcriptomic data. As one of these initiatives, we introduced ArrayAnalysis, in 2013 as a web-based application designed as a user-friendly solution for microarray data analysis [1]. While this earlier version helped many researchers analyse microarray experiments more easily, it did not support RNA-seq data and lacked more advanced interactive features. More recently, other applications have been developed for the analysis of RNA-seq data [2–5].

Although these applications lower the barrier to entry, they still have important limitations. Many provide little guidance to help users navigate the analysis workflow, making them less suitable for researchers inexperienced with gene expression profiling. In addition, the output of such tools is often rigid, with limited options to support interactive data exploration and to customise figures for publication-quality presentation. To address these limitations, we developed a new version of ArrayAnalysis, a web-based application that provides a streamlined and accessible workflow for the analysis of both microarray and RNA-seq data. ArrayAnalysis offers several innovations, including enhanced user guidance throughout the analysis steps, interactive outputs that facilitate data exploration, and customisable figures tailored for use in scientific publications. With these features, ArrayAnalysis aims to make transcriptomic data analysis more approachable and effective for a broad audience of biomedical researchers.

## IMPLEMENTATION

We developed ArrayAnalysis (https://arrayanalysis.org) as a tool for user-friendly analysis of transcriptomic data. The platform is built with the Shiny framework in the R programming language (http://www.r-project.org) and integrates more than 30 packages from CRAN [6–28] and Bioconductor [29–43] (**Table 1**). ArrayAnalysis is made available as a web application and for local installation as a desktop app, Docker image, and R package.

**Table 1.**
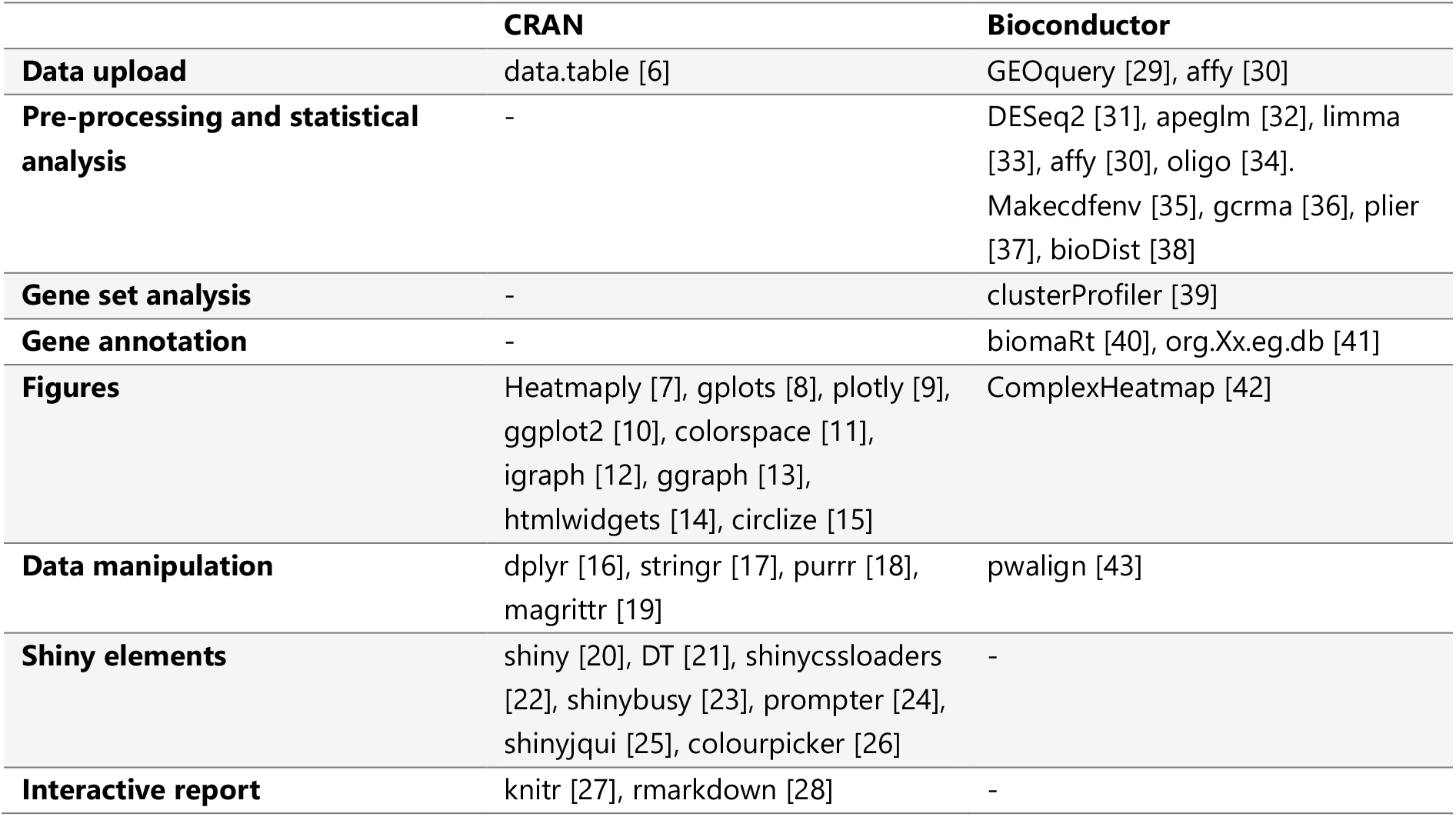
CRAN and Bioconductor packages used by ArrayAnalysis.

### Input

In the first step of the ArrayAnalysis app, users can upload expression data and associated metadata. Microarray data can be provided as a zipped folder containing CEL files, as an expression matrix (.tsv/.csv/.txt), or as a GEO series matrix file. RNA-seq data can be uploaded as a matrix of raw or normalized counts (.tsv/.csv/.txt). The metadata, which contains information about the samples such as the experimental group and age, can be provided as a table (.tsv/.csv/.txt) or as a GEO series matrix file. Upon uploading the data, a preview of both the expression data and metadata is displayed, allowing users to verify that the files have been correctly loaded. If necessary, the metadata can be modified within the application.

ArrayAnalysis includes two demo datasets from the Gene Expression Omnibus (GEO) [44] that can be directly loaded within the app: GSE6955 [45] (microarray data) and GSE128380 [46] (RNA-seq data). Both datasets comprise gene expression profiles of post-mortem brain tissue from Rett syndrome patients and age-matched controls. The GSE6955 and GSE128380 datasets use the Affymetrix Human Genome U95 v2 array and Illumina HiSeq 2000 platforms, respectively. The GSE128380 dataset is used to generate the figures in this manuscript.

### Output

Following the data upload, users can perform pre-processing and quality control, differential gene expression analysis, and gene set analysis. To ensure users follow the correct workflow, modules become accessible sequentially—each step must be completed before proceeding to the next. Additionally, to guide users through the analysis, each module includes explanations of the analysis settings (*e*.*g*., **Figure 1A**) and of all provided figures and tables. ArrayAnalysis is designed to generate outputs that are 1) interactive and facilitate data exploration as well as 2) downloadable as publication-ready figures. The latter is achieved by making all figures customisable, including options for changing the layout and colour palettes (**Figure 1B**). Figures can be exported as vector graphics (PDF) or high-resolution raster images (PNG, TIF at 300 dpi). Interactive plots can also be saved as a standalone HTML file. Finally, each module offers the option to download an interactive report summarising the analysis.

**Figure 1.**
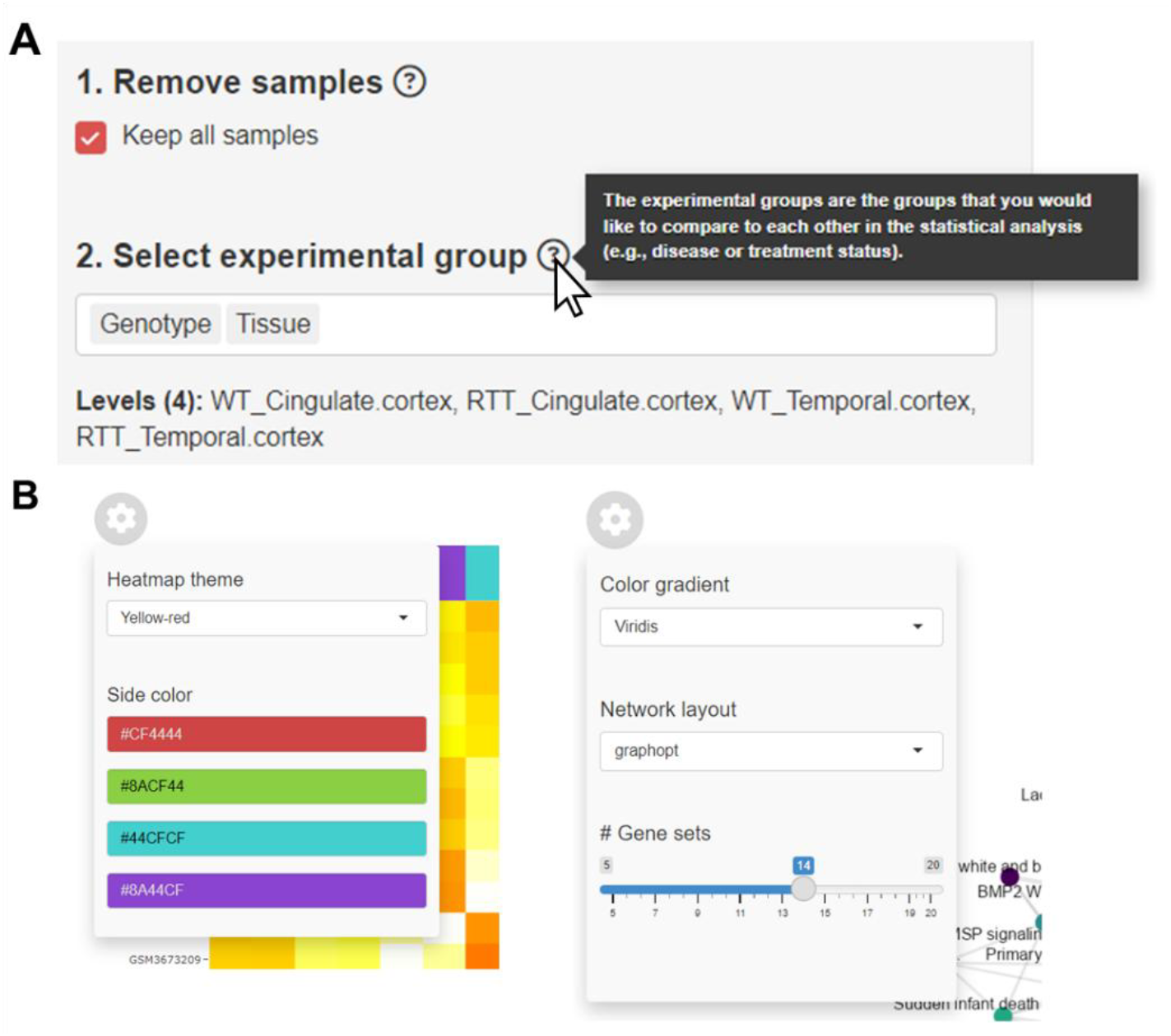
User-guidance and figure customisation in ArrayAnalysis. **A)** Each module includes in-app information boxes with explanations of the analysis settings. **B)** Each figure can be interactively customised by the user.

## RESULTS

We propose ArrayAnalysis as a user-friendly solution for transcriptomic data analysis. Following data upload, users can analyse their data in three sequential steps: 1) pre-processing and quality control, 2) differential gene expression analysis, and 3) gene set analysis. In the following sections, we provide a detailed description of these three functionalities and perform a comparison with other existing transcriptomics data analysis tools.

### Pre-processing and quality control

In the pre-processing and quality control module, the user can prepare the data for downstream analysis, including interactive filtering of genes and samples as well as normalization and transformation of expression data. In addition, the data are automatically screened for potential outlying samples. If potential outliers are detected, a warning message is displayed to alert the user. An outlier warning is provided when there is at least one sample that deviates by more than six median absolute deviations from the median on at least one principal component that explains ≥10% of the total variance.

Several quality control outputs are generated to assist users in evaluating the data quality and identifying potential outliers. These include boxplots of gene expression distributions, density plots, sample–sample correlation heatmaps, and principal component analysis (PCA) plots (**Figure 2**). Furthermore, an interactive gene expression matrix is provided that allows users to query specific genes and visualize their expression patterns across experimental groups. For analyses starting from a raw RNA-seq count matrix, an additional bar chart depicting total read counts per sample is included to help identify discrepancies in sequencing depth.

**Figure 2.**
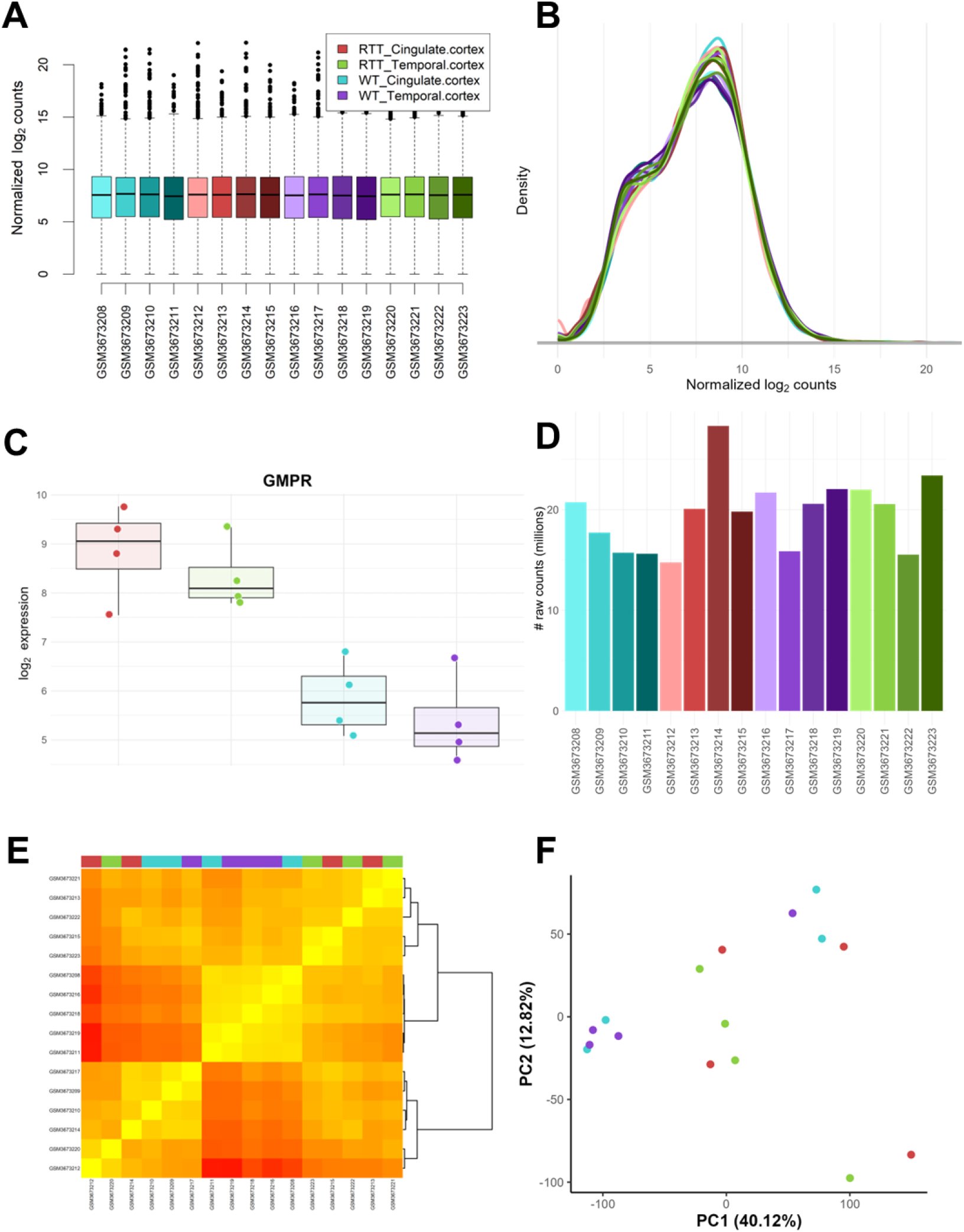
Quality control plots provided by ArrayAnalysis. **A)** Sample-wise expression boxplot. **B)** Sample-wise density plot. **C)** Expression boxplot of a user-selected gene. **D)** Bar chart of total raw read counts; this plot is only provided when analysing raw RNA-seq count data. **E)** Heatmap of sample-sample correlations. **F)** PCA score plot. The density plot, heatmap, and PCA score plot are interactive and support zooming and on-hover data display. Detailed information about these plots can be found at https://arrayanalysis.org/explain.

### Differential gene expression analysis

In the differential gene expression analysis module, users can select the statistical comparison(s) of interest, select covariates, and map between gene identifier types. The *DESeq2* R/Bioconductor package [31] is used for the differential gene expression analysis of raw RNA-seq counts. For other input data, the *limma* R/Bioconductor package [33] is applied. The gene identifier mappings are extracted from the Ensembl database using the biomaRt R/Bioconductor package [40]. If the connection to the Ensembl database fails (e.g., due to server unavailability or no internet connection), gene mapping will be performing using the relevant Bioconductor annotation package (org.Xx.eg.db). The provided outputs include an interactive statistics table, histogram of p-values and log_2_ fold changes (log_2_FCs), and interactive volcano and MA plots (**Figure 3**).

**Figure 3.**
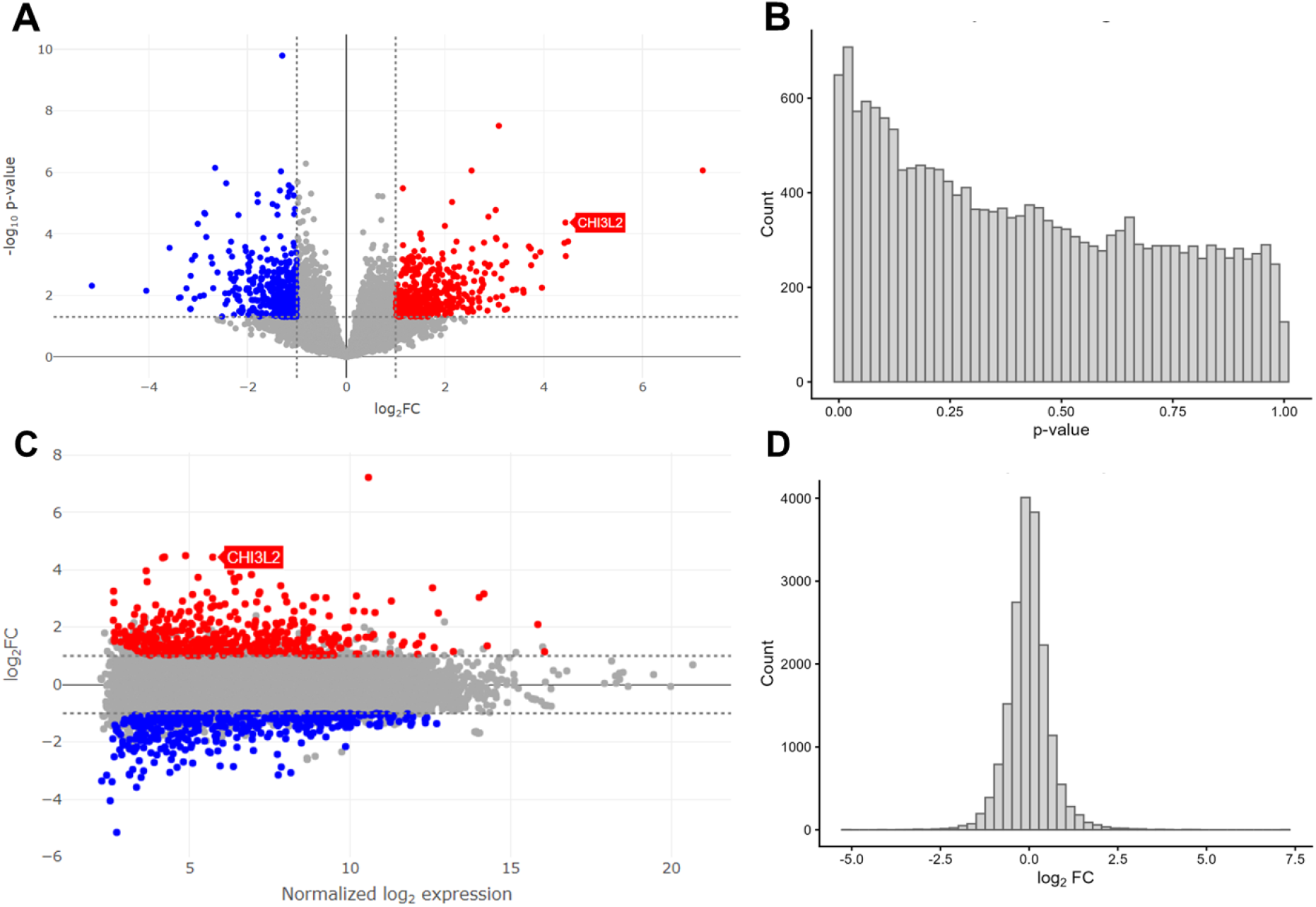
Differential gene expression analysis plots provided by ArrayAnalysis. **A)** Volcano plot. **B)** p-value histogram. **C)** MA plot. **D)** log_2_FC histogram. All four plots have zooming and on-hover data display functionalities. Detailed information about these plots can be found at https://arrayanalysis.org/explain.

### Gene set analysis

Following the differential gene expression analysis, ArrayAnalysis uses the *clusterProfiler* R/Bioconductor package [39] to support two types of gene set analysis: overrepresentation analysis and gene set enrichment analysis. These analyses can be performed on gene ontologies (GO-BP, GO-MF, and GO-CC) [47], KEGG [48], and WikiPathways [49] gene sets. In this module, an interactive statistics table, bar chart, and gene set similarity network are provided (**Figure 4**).

**Figure 4.**
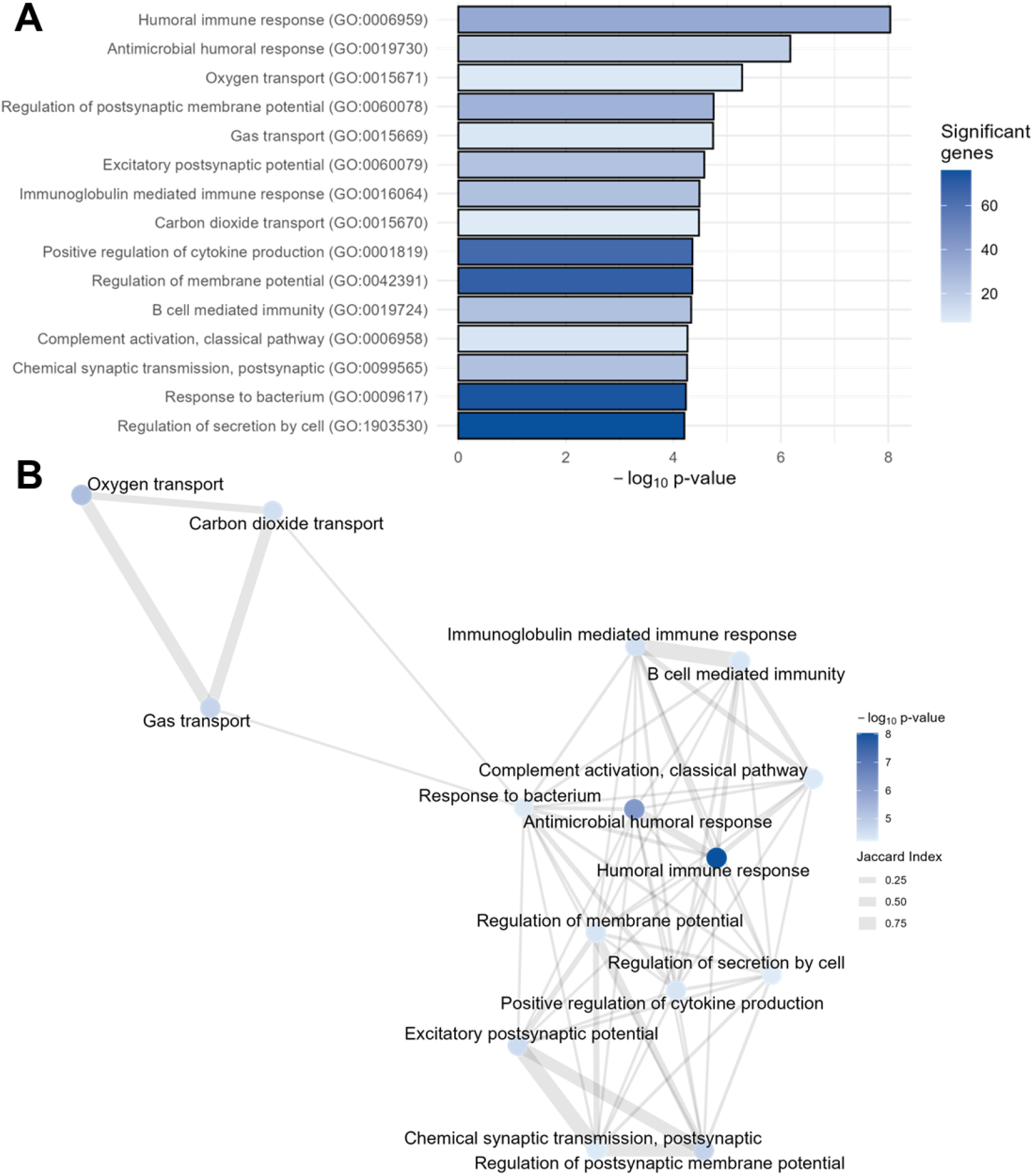
Gene set analysis plots provided by ArrayAnalysis. **A)** Interactive gene set analysis bar chart. **B)** Gene set similarity network. Detailed information about these plots can be found at https://arrayanalysis.org/explain.

### Comparison with other tools

We compared ArrayAnalysis with other existing web applications for transcriptomic analysis [2–5] (**Table 2**). Relative to these tools, the new version of ArrayAnalysis offers a broader range of options for local installation and supports a more diverse set of input data. Furthermore, the application has several unique features, including automatic outlier detection, in-app sample removal and metadata editing, and customisation of all figures. Finally, compared to other web applications, ArrayAnalysis provides a wider range of figure export formats.

**Table 2.** Comparison of ArrayAnalysis with other web applications for transcriptomic analysis.

|  | **ArrayAnalysis (2026)** | **STAGEs (2023)** | **GeneCloud-Omics (2021)** | **GENAVi (2019)** | **iDEP (2018)** | **ArrayAnalysis (2013)** |
| --- | --- | --- | --- | --- | --- | --- |
| **Availability** | Web app, Docker image, desktop app, R package | Web app, Docker image | Web app, Docker image | Web app, Docker image | Web app, Docker image, R package | Web app |
| **Accepted microarray data** | CEL files, raw and normalized intensity table | Normalized intensity table | CEL files | N/A | Normalized intensity table | CEL files, intensity table |
| **Accepted RNA-seq data** | Raw and normalized count table | Raw and normalized count table | Raw and normalized count table | Raw count table | Raw and normalized count table | N/A |
| **Automated outlier detection** | Yes | No | No | No | No | No |
| **In-app sample removal** | Yes | No | No | No | No | No |
| **In-app metadata editing** | Yes | No | No | No | No | No |
| **Figure customisation** | Yes | No | No | No | Some figures | No |
| **Figure export format** | Vector (PDF), raster (PNG/TIF), and interactive (HTML) | Raster (PNG, < 300 dpi) | Vector (PDF) and raster (PNG) | Raster (PNG, < 300 dpi) | Vector (PDF/SVG) and raster (PNG) | Vector (PDF) and raster (PNG) |

## DISCUSSION

We developed a new version of ArrayAnalysis (https://arrayanalysis.org) as a user-friendly solution for transcriptomic data analysis. Compared to the first release in 2013 [1], the current version has been substantially updated; besides offering support for RNA-seq data, it introduces a strongly interactive analysis workflow that facilitates efficient data exploration and analysis. It further addresses key limitations of existing tools by providing extensive user-guidance and customisable, publication-ready outputs. Finally, in addition to the web application, ArrayAnalysis can be run locally as an R package, desktop application, and Docker image, enabling secure in-house processing of sensitive data.

A central aim of ArrayAnalysis is to support a broad community of users, ranging from biomedical researchers without prior bioinformatics training to experienced data analysts. For users with less bioinformatics experience, the platform offers elaborate guidance features, including in-app information boxes, tutorials, FAQs, and extensive documentation. In addition, automated outlier detection helps users recognize potential issues in their data.

For expert users, ArrayAnalysis provides a streamlined and flexible workflow. Options such as sample exclusion, metadata editing, and figure customisation can be applied directly within the application, reducing the need to alternate between multiple tools. The strength of ArrayAnalysis lies in providing a streamlined and robust workflow for core transcriptomic data analysis functions. Moreover, to support the use of more specialized tools for advanced downstream tasks, results can be downloaded at each step and are designed to ensure interoperability with other applications. For example, differential expression statistics are provided in a consistent tabular format across different input data types and include standard error estimates of log_2_ fold changes, enabling integration into subsequent analytical pipelines. Moreover, the semi-automatic and interactive workflow enables users to perform core transcriptomic analysis tasks reliably within very limited time, including the detection and removal of outliers without the need to re-upload data. These features make ArrayAnalysis particularly well-suited for systematic (re)processing or large-scale, transcriptome-wide meta-analyses that require efficient and consistent processing of dozens of datasets.

The ability to generate publication-ready visualizations is a core feature that benefits both novice and experienced bioinformatics researchers. Figures can be customised in terms of colour palettes and various other layout elements as well as exported in different aspect ratios and file formats, including high-resolution raster images and vector graphics that meet journal requirements. The option to export session information and user-selected analysis settings ensures transparency and reproducibility, as well as aiding manuscript preparation.

## CONCLUSIONS

In conclusion, ArrayAnalysis provides both wet-lab scientists and bioinformaticians with a powerful, intuitive, and freely available tool for transcriptomic data analysis. By lowering barriers to entry while also supporting expert-level workflows, ArrayAnalysis contributes to more widespread, reproducible, and effective use of transcriptomics in biomedical research.

## AVAILABILITY AND REQUIREMENTS

**Project name:** ArrayAnalysis

**Project home page:** https://arrayanalysis.org/

**Operating system(s):** Platform independent

**Programming language:** R

**License:** CC-BY-4.0

**Any restrictions to use by non-academics:** None

## LIST OF ABBREVIATIONS

RNA-seq: RNA sequencing
log_2_FC: log_2_ fold change

## DECLARATIONS

### Ethics approval and consent to participate

Not applicable.

### Consent for publication

Not applicable.

### Availability of data and materials

ArrayAnalysis is freely available for all users at https://arrayanalysis.org under the Creative Commons Attribution 4.0 International (CC BY 4.0) license. The source code is available at https://doi.org/10.5281/zenodo.19957055.

### Competing interests

Not applicable.

### Funding

This work was supported by Stichting Terre – The Dutch Rett Syndrome foundation and the Virtual Human Platform for Safety Assessment project, which is funded by the Netherlands Research Council (NWO) ‘Netherlands Research Agenda: Research on Routes by Consortia’ [NWA-ORC 1292.19.272]

### Authors’ contributions

Jarno Koetsier: Conceptualization, Software, Project administration, Visualization, Writing— original draft. Ozan Cinar: Software, Writing—review & editing. Egon L. Willighagen: Resources, Writing—review & editing. Ammar Ammar: Software, Writing—review & editing. Vishnu Karthik: Software, Writing—review & editing. Denyel Jennen: Validation, Writing—review & editing. Chris T. Evelo: Writing—review & editing. Leopold M.G. Curfs: Funding acquisition, Writing—review & editing. Chris P. Reutelingsperger: Funding acquisition, Writing—review & editing. Nasim Bahram Sangani: Resources, Writing—review & editing. Lars M.T. Eijssen: Conceptualization, Supervision, Project administration, Writing—original draft.

## Acknowledgements

Not applicable.

## Notes

https://arrayanalysis.org

### Competing Interest Statement

The authors have declared no competing interest.

